# Riboflavin Deficiency-Induced Angular Stomatitis and Conjunctivitis in Associated with Gut Microbiota Dysbiosis

**DOI:** 10.64898/2026.01.30.702774

**Authors:** Wei Zhang, Pan-pan Wang, Wen-qi Shi, Hui Zhao, Feng Pan

## Abstract

**Purpose:** Riboflavin deficiency causes ariboflavinosis, and the purpose of this study is to investigate the potential biological factors underlying its occurrence.

**Methods:** Male F344 rats were randomly assigned to R6 and R0 groups. The healthy controls (R6 group) was fed a riboflavin-normal diet (6 mg/kg riboflavin), while the R0 group was fed a riboflavin-deficient diet (0 mg/kg riboflavin) for 16 weeks. Following this period, the R0 group was switched to a riboflavin replenishment diet (6 mg/kg riboflavin) for an additional 4 weeks (weeks 17-20). The bacterial communities were analyzed using high-throughput 16S rRNA gene sequencing.

**Results:** Riboflavin deficiency induces ariboflavinosis in rats (66.7%), characterized by angular stomatitis and conjunctivitis. With riboflavin replenishment, ariboflavinosis was completely resolved. Riboflavin deficiency altered the gut microbiota co-occurrence network and composition. The potential mechanism is predicted to involve an increase in glycan biosynthesis and metabolism within the gut microbiota, particularly in lipopolysaccharide biosynthesis pathways.

**Conclusions:** These findings suggest that riboflavin deficiency induces ariboflavinosis by altering the gut microbiota, providing new insights into the mechanisms of riboflavin deficiency and its association with chronic inflammation.

**Clinical perspectives:** Riboflavin is an essential nutrient that must be obtained through the diet, as the human body cannot synthesize it. Since riboflavin is not stored in the body, regular dietary intake is necessary. Riboflavin deficiency is prevalent worldwide and can lead to ariboflavinosis. However, the specific mechanisms underlying riboflavin deficiency-induced ariboflavinosis remain unclear. This study demonstrates that riboflavin deficiency causes ariboflavinosis (characterized by angular stomatitis and conjunctivitis) in rats and reveals a close relationship between riboflavin deficiency and alterations in gut microbiota composition. Gut dysbiosis appears to be a hallmark of ariboflavinosis and may serve as a promising diagnostic biomarker and therapeutic target for the condition.

## Background

Riboflavin (vitamin B2) is a water-soluble vitamin found in a wide variety of foods. As the human body cannot synthesize riboflavin, it is essential to include it in the diet, requiring daily replenishment through food consumption^[1]^. Good dietary sources of riboflavin include dairy products, meats, eggs, fish, and certain fruits and dark-green vegetables. However, according to the recommendations of the American Food and Nutrition Board (1.3 mg/day), a significant portion of the population does not meet the daily nutritional requirement for this vitamin^[2]^. While certain microorganisms in the human intestine can produce riboflavin, their concentrations are insufficient to prevent deficiency^[3]^.

Riboflavin deficiency is widespread in many regions around the world, particularly in underdeveloped countries where there is a low intake of dairy products and meats ^[4, 5]^. This deficiency also commonly among pregnant women, lactating women, infants, schoolchildren, and the elderly ^[1, 6, 7]^. Riboflavin deficiency causes ariboflavinosis, a common nutritional deficiency disorder characterized by a range of clinical abnormalities, including sore throat, glossitis (inflammation of the tongue), angular stomatitis (cracks at the corners of the mouth), and seborrheic dermatitis^[8, 9]^.

Diet plays a pivotal role in shaping the composition of the gut microbiome, with various diets having a profound impact on the stability, functionality, and diversity of the microbial community within our gut^[10]^. The gut microbiome is intricately involved in multiple aspects of host physiology throughout life, from the development and maturation of immune responses to stress response and behavior^[11]^. Understanding the profound impact of varied diets on the microbiome is crucial, as it will enable us to make well-informed dietary decisions for better metabolic health and help prevent and slow the onset of specific diet-related diseases. We hypothesize that dietary riboflavin deficiency will affect the gut microbiota. The relationship between the gut microbiota and riboflavin deficiency-induced disease remains unknown.

In the present study, we investigate the effects of riboflavin deficiency on ariboflavinosis in F344 rats. To elucidate the underlying mechanisms, we analyze the composition of the gut microbiota using 16S rRNA gene amplicon sequencing. Additionally, we study the correlation between the gut microbiota and riboflavin deficiency-induced disease.

## Materials and Methods

### Animal experiments

Male F344 rats (SPF, 5-week-old) were purchased from the Beijing Vital River Laboratory Animal Technology Co., Ltd. Rats were allowed to acclimate for 1 week in a controlled environment (temperature 23±2 °C and humidity 50±5%, with a 12-hour light-dark cycle). The experimental flow chart is shown in **Figure 1A**. The rats (*n* = 20) were randomly assigned to two groups. The healthy controls (R6 group) was fed a riboflavin-normal diet (AIN-93M purified diet, containing 6 mg/kg riboflavin, Nantong Trophic Animal Feed High-Tech Co., Ltd., China)^[12]^. The R0 group was fed a riboflavin-deficient diet (made with the same formula, but containing 0 mg/kg riboflavin) for weeks 0-16, and then switched to a diet containing 6 mg/kg riboflavin for weeks 17-20. The experiment lasted for 20 weeks. Rat ariboflavinosis-free survival was closely monitored throughout the experimental period. At weeks 5, 10, 16, and 20, fecal samples were collected in sterile Eppendorf tubes and immediately stored at -80°C for subsequent analysis. At the end of the experiment, blood samples were collected from the orbital plexus after pentobarbital sodium anesthesia. The rats were then euthanized using the carbon dioxide suffocation method.

**Figure 1.**
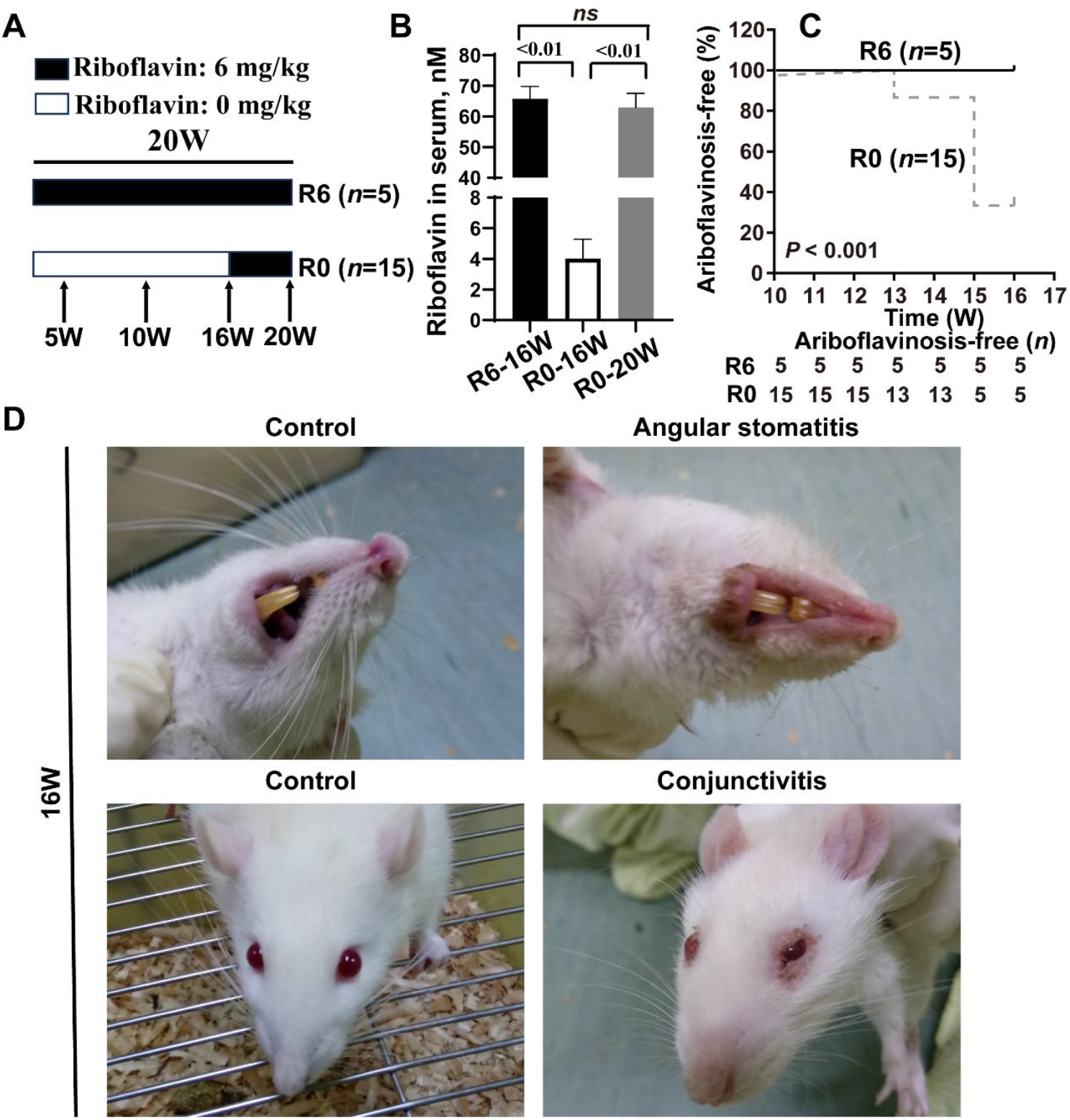
Riboflavin deficiency induces angular stomatitis and conjunctivitis in rats. **(A)** Schematic diagram showing the study design. (**B**) Riboflavin concentration in serum. (**C**) Kaplan-Meier curve illustrating the effect of riboflavin deficiency on ariboflavinosis-free survival. (**D**) Representative pictures of angular stomatitis, and conjunctivitis. Data are means ± SD, *n* = 5-15/group, differences were analyzed using an unpaired Student’s *t*-test.

### Measurement of riboflavin

The concentration of riboflavin in serum was measured by high performance liquid chromatography (HPLC, Agilent 1200 system) as described previously ^[13]^. Briefly, blood samples were centrifuged at 1,500 g for 15 minutes to obtain serum. Proteins were removed from the serum by acid precipitation, and an aliquot of the resulting supernatant was analyzed by HPLC. Impurities were separated from riboflavin isocratically, and the target material was detected fluorometrically (excitation 450 nm; emission 520 nm).

### Gut Microbiota Analysis

5-10 rats were selected from R6 and R0 group (rats that exhibit symptoms of angular stomatitis and conjunctivitis) for gut microbiota analysis. The cage as a variable in the final statistical analyses^[14]^. Total bacterial DNA was extracted from fecal samples using the CTAB/SDS method and then checked by agarose gel electrophoresis ^[15]^. The bacterial 16S rRNA gene encompassing the V3-V4 hypervariable regions was amplified by PCR using barcoded primers adapted from existing 341F and 806R universal primers. Sequencing was performed on an Illumina MiSeq PE250 platform at Biozeron Biological Technology Co, Ltd (Shanghai, China). Paired-end reads (250bp/300bp) from the original DNA fragments were merged using FLASH (v1.2.11) ^[16]^. The obtained reads were chimera checked compared with the Gold Database (Release v7) and clustered into operational taxonomic units (OTUs) by UPARSE software (version 7.1, http://drive5.com/uparse/) with a 97% similarity cutoff. Representative sequences was selected for each OUT, and the RDP classifier was used to annotate taxonomic information for each representative sequence. In order to compute Alpha Diversity, the OTU table was rarified, and calculate three metrics: Chao1 estimates the species abundance; Observed Species estimates the amount of unique OTUs found in each sample, and Shannon index. Rarefaction curves were generated based on these three metrics. Unweighted unifrac distance for Principal Coordinate Analysis (PCoA) was performed with QIIME2 (v2019.4) ^[17]^. To confirm differences in the abundances of individual taxa between the two groups, LEfSe analysis was utilized. A Kruskal-Wallis sum-rank test was performed to examine the changes and dissimilarities among classes, followed by Linear Discriminant Analysis (LDA) analysis to determine the contribution effect of each distinctively abundant taxa. To identify differences in microbial communities between the two groups, ANOSIM (Analysis of Similarity) and MRPP (multi-response permutation procedure) were performed based on the Bray-Curtis dissimilarity distance matrices. Microbial community functions were predicted by PICRUSt2 Software^[18]^, and statistics analysis and plotting were conducted with STAMP Software ^[19]^.

### Statistical Analysis

Statistical analysis was performed using SPSS 22.0 software (SPSS, Chicago, IL). Data are presented as means ± SD. The unpaired Student’s *t*-test (unequal variances, Mann-Whitney test) was used to compare the two groups. Statistical tests were two-sided, and significance was set at *P* < 0.05.

## Results

### Riboflavin deficiency induces angular stomatitis and conjunctivitis in rats

At 16 weeks, the riboflavin concentration in serum was significantly lower in the riboflavin-deficient group (R0) compared to the riboflavin-normal group (R6) (**Figure 1B**, *P* < 0.001). Following riboflavin replenishment, the serum riboflavin concentration in the R0 group returned to the level of R6 group at 20 weeks (*P* > 0.05, **Figure 1B**). From the 13th week, ariboflavinosis occurred exclusively in the R0 rats, characterized by angular stomatitis and conjunctivitis, which appeared either simultaneously or alone (angular stomatitis=6 rats, angular stomatitis and conjunctivitis =4 rats). Ariboflavinosis occurred in 66.7% (10 out of 15) of rats in the R0 group, whereas all rats (100%) in the R6 group were remained normal (**Figure 1C**). Representative pictures of angular stomatitis and conjunctivitis are shown in **Figure 1D**. With riboflavin replenishment for 4 weeks, ariboflavinosis was completely resolved (100%). Taken together, riboflavin deficiency induces ariboflavinosis, and riboflavin replenishment is effective in treating this condition.

### Riboflavin deficiency modulates gut microbiota diversity and co-occurrence network in rats

The trend of individual rarefaction curves (**Supplemental Figure 1A**) and Shannon-Wiener curves (**Supplemental Figure 1B**) indicated that the high sampling coverage was achieved in all samples. As shown in **Figure 2A**, there was no significant difference in bacterial richness (Chao1 index) and bacterial diversity (Shannon index) between R0-16W rats and the R6-16W rats (*P* > 0.05). Riboflavin replenishment, the bacterial richness (Chao1 index) of the R0-20W rats increased compared with the R0-16W rats ((**Figure 2B**, *P* = 0.003); however, there was no significant difference in bacterial diversity (Shannon index). Principal coordinates analysis (PCoA) revealed a clear separation between the gut microbiota of the R0-16W rats and R6-16W rats (**Figure 2C**); riboflavin replenishment, there is still a noticeable separation between the gut microbiota of the R0-20W rats and R0-16W rats (**Figure 2D**). This indicates that the riboflavin-deficient diet had a substantial effect on gut microbiota composition. Analysis of relative abundance at the phylum level showed that the abundance of Proteobacteria was higher in the R0-16W rats compared to the R6-16W rats (**Figure 2E)**. Riboflavin deficiency or replenishment affected the abundance of Proteobacteria (**Figure 2F)**. The co-occurrence network of gut microbiota shows that Firmicutes and Bacteroidetes were co-occurrence in R6-16W rats (**Figure 3A-C)**. In contrast, R0-16W rats showed co-occurrence of Firmicutes, Bacteroidetes and Proteobacteria (**Figure 3D-F)**. Collectively, these results indicate that the gut microbiota is modulated by riboflavin deficiency in rats.

**Figure 2.**
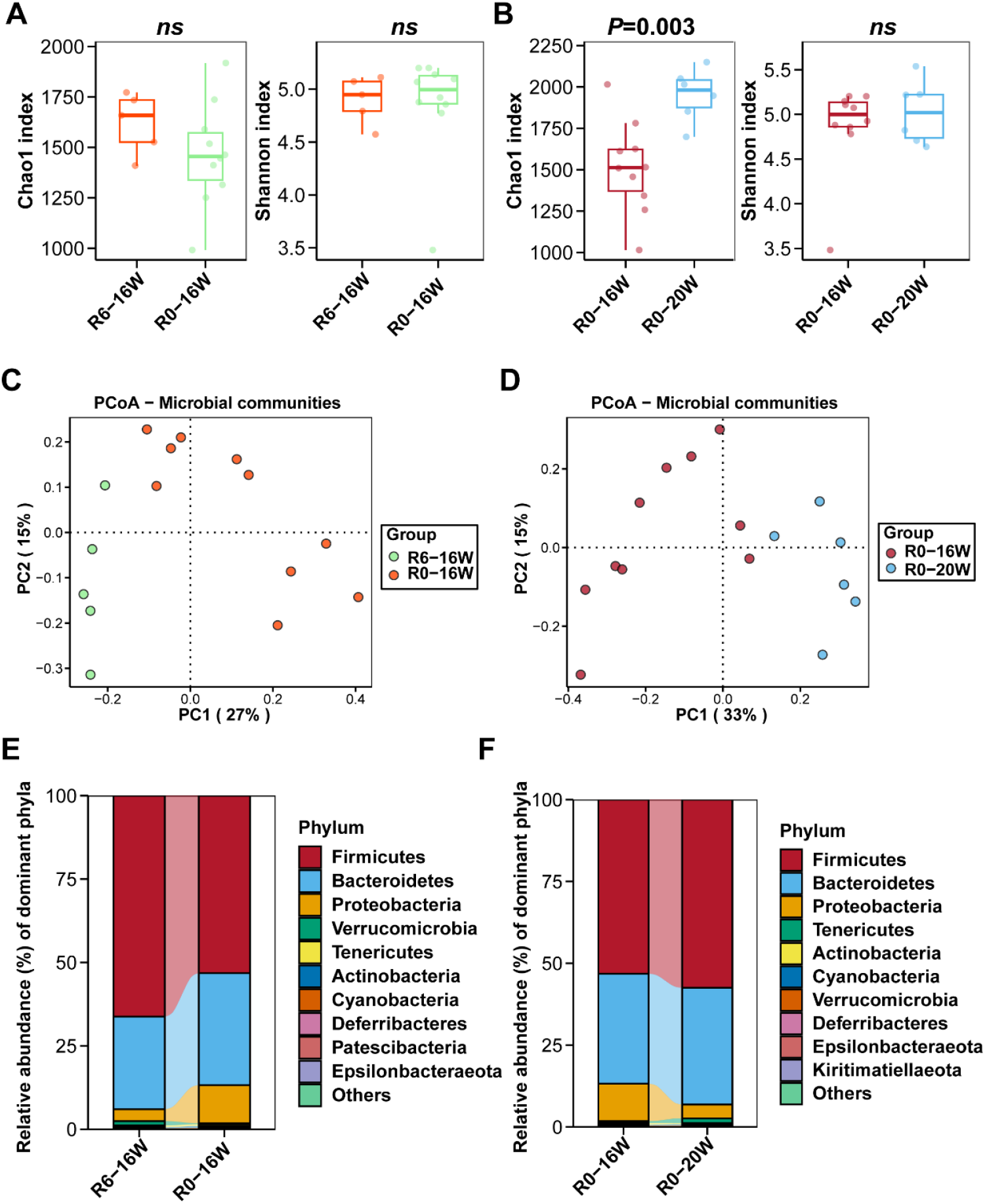
Riboflavin deficiency modulates gut microbiota diversity and composition at the phyla level in rats. Chao1 richness estimator and Shannon diversity index of gut microbiota in riboflavin deficiency (**A**) and riboflavin replenishment (**B**). Principal coordinates analysis (PCoA) of gut microbiota based on OTU abundance in riboflavin deficiency (**C**) and after riboflavin replenishment (**D**). Gut microbiota composition at the phylum level in riboflavin deficiency (**E**) and after riboflavin replenishment (**F**).

**Figure 3.**
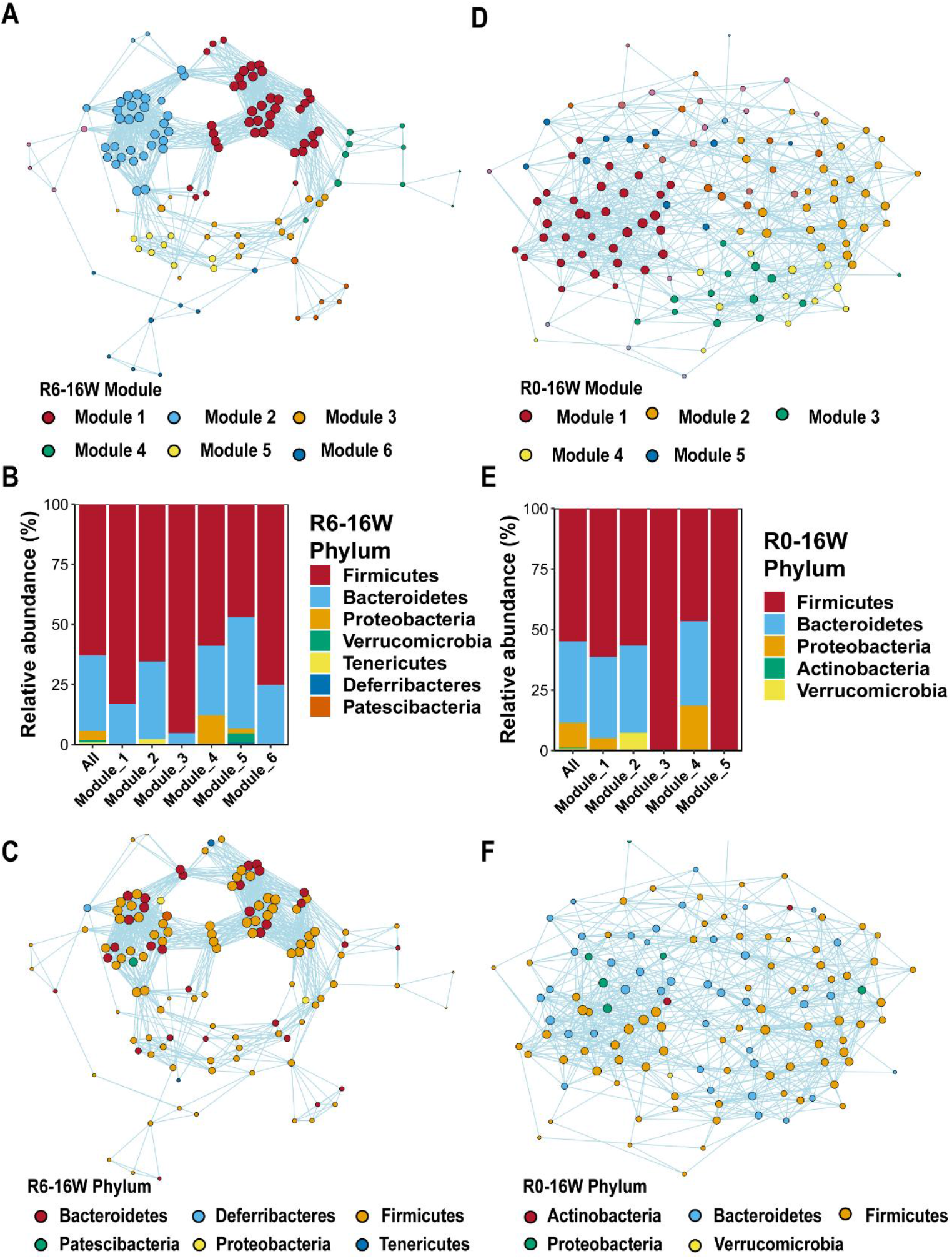
Riboflavin deficiency modulates gut microbiota co-occurrence network in rats. Riboflavin-normal, gut microbiota co-occurrence module (**A**), and module of gut microbiota composition (**B**). (**C**) Riboflavin-normal, gut microbiota co-occurrence network at the phylum level. (**D and E**) Riboflavin-deficient, gut microbiota co-occurrence module, and module of gut microbiota composition. (**F**) Riboflavin-deficient, gut microbiota co-occurrence network at the phylum level.

### Riboflavin deficiency modulates gut microbiota community and differential species in rats

The top 10 bacteria genera in the gut microbiota community of R0-16W rats are: *Lachnospiraceae NK4A136 group, Bacteroides, Ruminococcus 1, Ruminococcaceae UCG-005, Escherichia-Shigella, Turicibacter, Ruminiclostridium 9, Christensenellaceae R-7 group, Roseburia, Prevotellaceae Ga6A1 group*. The abundance of *Bacteroides, Ruminococcaceae UCG-005* and *Escherichia-Shigella* in R0-16W rats was higher than that in R6-16W rats (**Figure 4A**). The abundance of *Bacteroides* and *Escherichia-Shigella* in R0-16W rats increased over time (5 weeks, 10 weeks, 16 weeks) (**Figure 4B**). With riboflavin replenishment for 4 weeks, the abundance of *Bacteroides, Ruminococcaceae UCG-005* and *Escherichia-Shigella* decreased in R0-20W rats (**Figure 4C**). In terms of the phylum level of gut microbiota, *Bacteroides* belong to the Bacteroidetes, *Escherichia-Shigella* belong to the Proteobacteria, and *Ruminococcaceae UCG-005* belong to the Firmicutes (**Figure 4D**). Analysis of relative abundance at the genus level showed that the abundance of *Bacteroides, Ruminococcaceae UCG-005* and *Escherichia-Shigella* in R0-16W rats was significantly higher than that in R6-16W rats (**Figure 4E)**.

**Figure 4.**
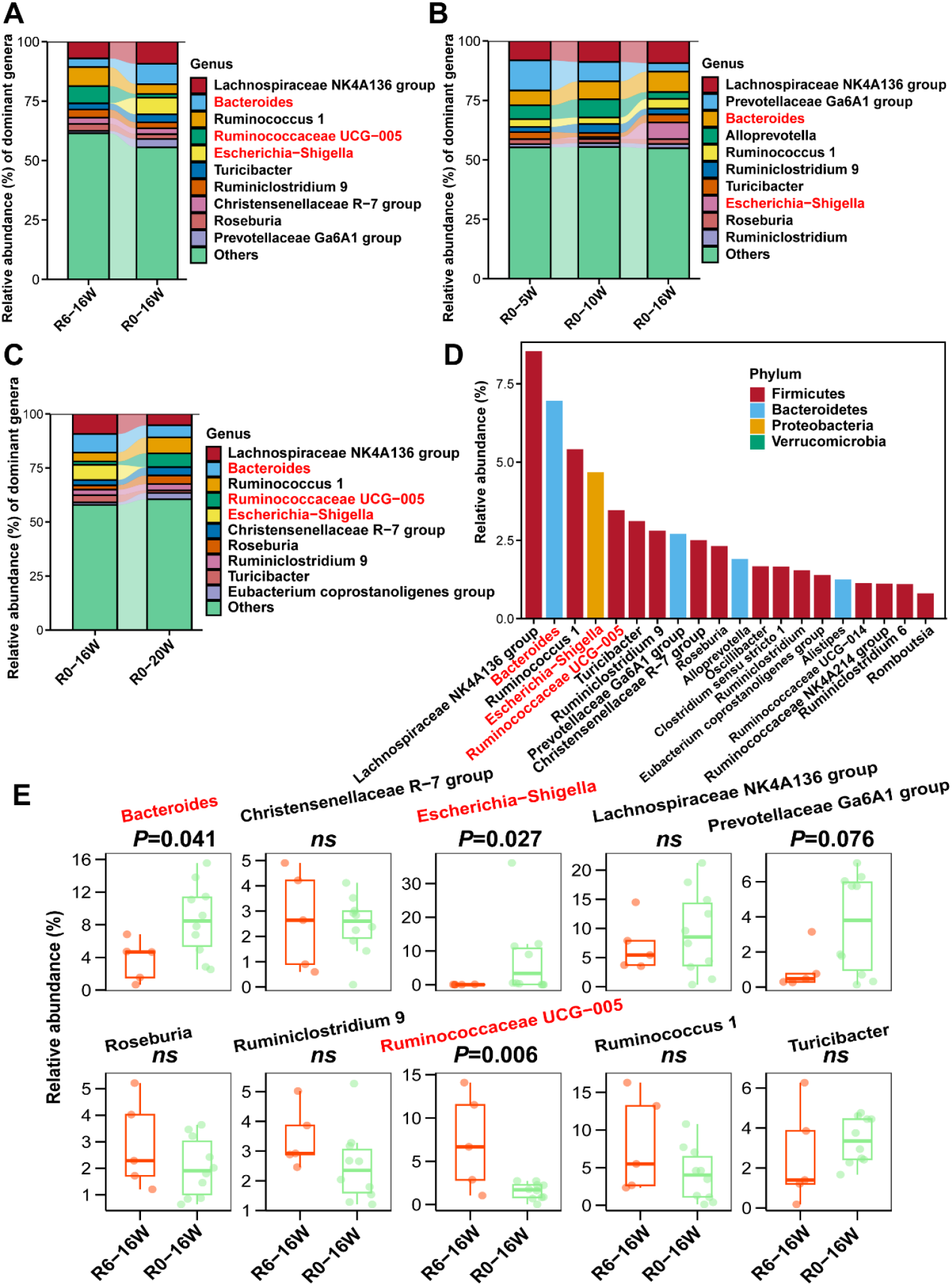
Riboflavin deficiency modulates gut microbiota community at the genus level in rats. (**A**) Riboflavin deficiency, the abundance of top 10 bacteria genera in the gut microbiota community. (**B**) Changes in the abundance of top 10 bacteria genera at different time points. (**C**) Riboflavin replenishment, the abundance of the top 10 bacteria genera in the gut microbiota community. (**D**) The corresponding phylum of the dominant genera. (**E**) Riboflavin deficiency causes the abundance differences of gut microbiota at the genus level.

The cladogram from LEfSe analysis indicates that bacterial markers distinguishing R0-16W rats include Actinobacteria (Bifidobacteriaceae, Bifidobacteriales), Bacteroidetes (Bacteroidaceae, Marinifilaceae, Prevotellaceae, Rikenellaceae, Tannerellaceae), Firmicutes (Clostridiales_vadinBB60_group), Proteobacteria (Enterobacteriaceae, Enterobacteriales, Gammaproteobacteria) (**Figure 5A)**. LDA analysis showed that Proteobacteria, specifically *Escherichia_Shigella*, Enterobacteriaceae, Enterobacteriales, Gammaproteobacteria, contributed the most to the significant differences in the community between R0-16W rats and R6-16W rats (**Figure 5B)**. Collectively, these results indicate that the gut microbiota community and differential species is modulated by riboflavin deficiency in rats.

**Figure 5.**
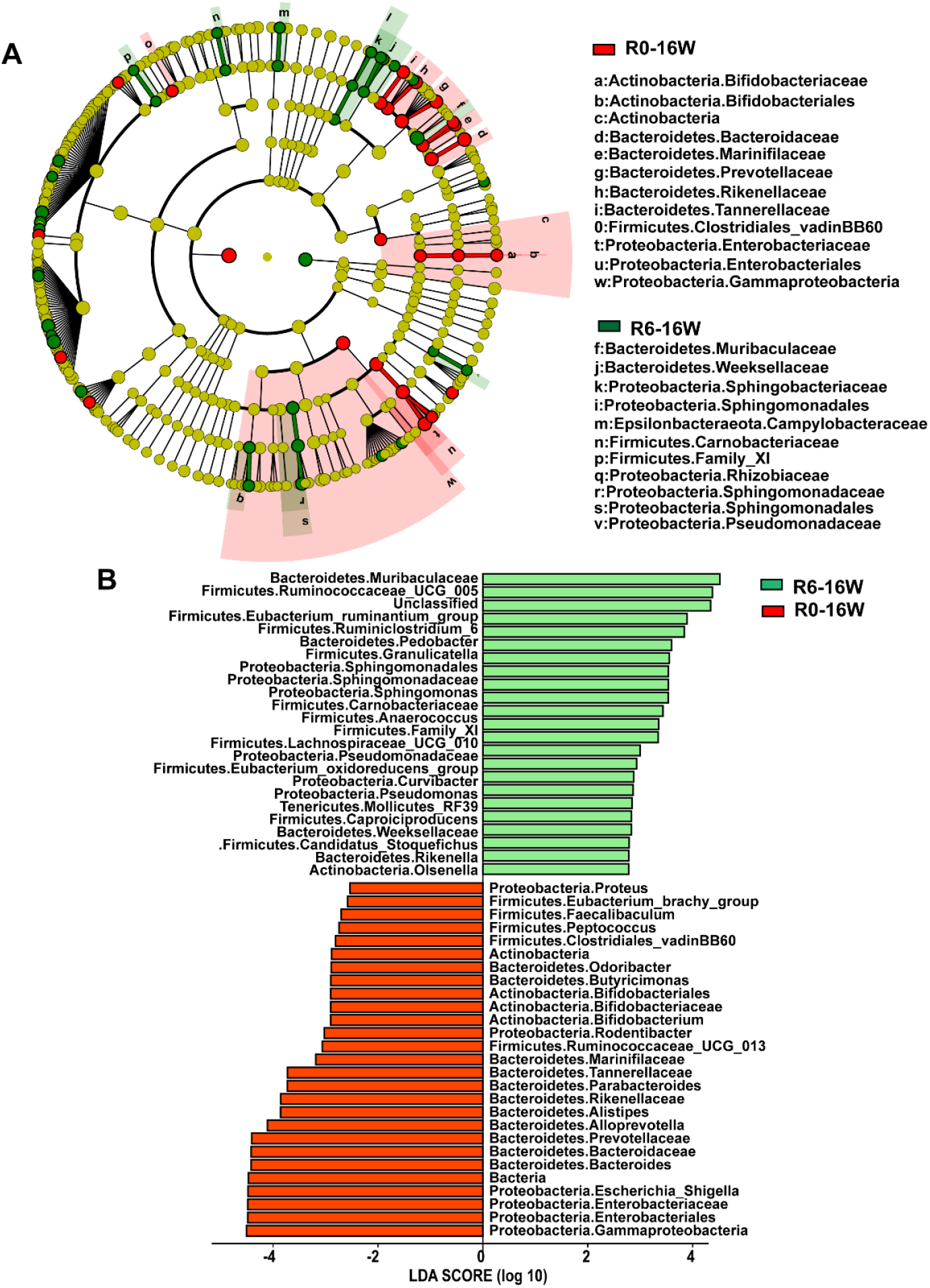
Cladogram and LDA analysis showing bacterial markers contributing most between R0-16W and R6-16W rats. (**A**) The cladogram of LEfSe analysis showing bacterial markers. (**B**) LDA scores showing the bacterial that contribute most to the differences between R0-16W and R6-16W rats.

### Effects of riboflavin deficiency on the potential functions of gut microbiota in rats

The differences in gut microbiota functional profiles between R0-16W rats and R6-16W rats were predicted by PICRUSt2 (**Figure 6**). Compared with the R6-16W rats, at KEGG pathway level 2, Carbohydrate metabolism, Glycan biosynthesis and metabolism, and Metabolism of other amino acids in R0-16W rats were significantly elevated (**Figure 6A**, *P* < 0.05). Consistent with this, at KEGG pathway level 3, lipopolysaccharide biosynthesis, Other glycan degradation, fructose and mannose metabolism, Galactose metabolism, and Amino sugar and nucleotide sugar metabolism are significantly elevated (**Figure 6B**, *P* < 0.05). After 4 weeks of riboflavin replenishment, compared to the riboflavin-normal group (R6-20W), the absence of the above-mentioned abnormal metabolic pathways in the R0-20W group indicates that these metabolic pathways have been fully restored to normal levels (**Figure 7A-B**). Taken together, dietary riboflavin deficiency contributed to the functional differences in the gut microbiota.

**Figure 6.**
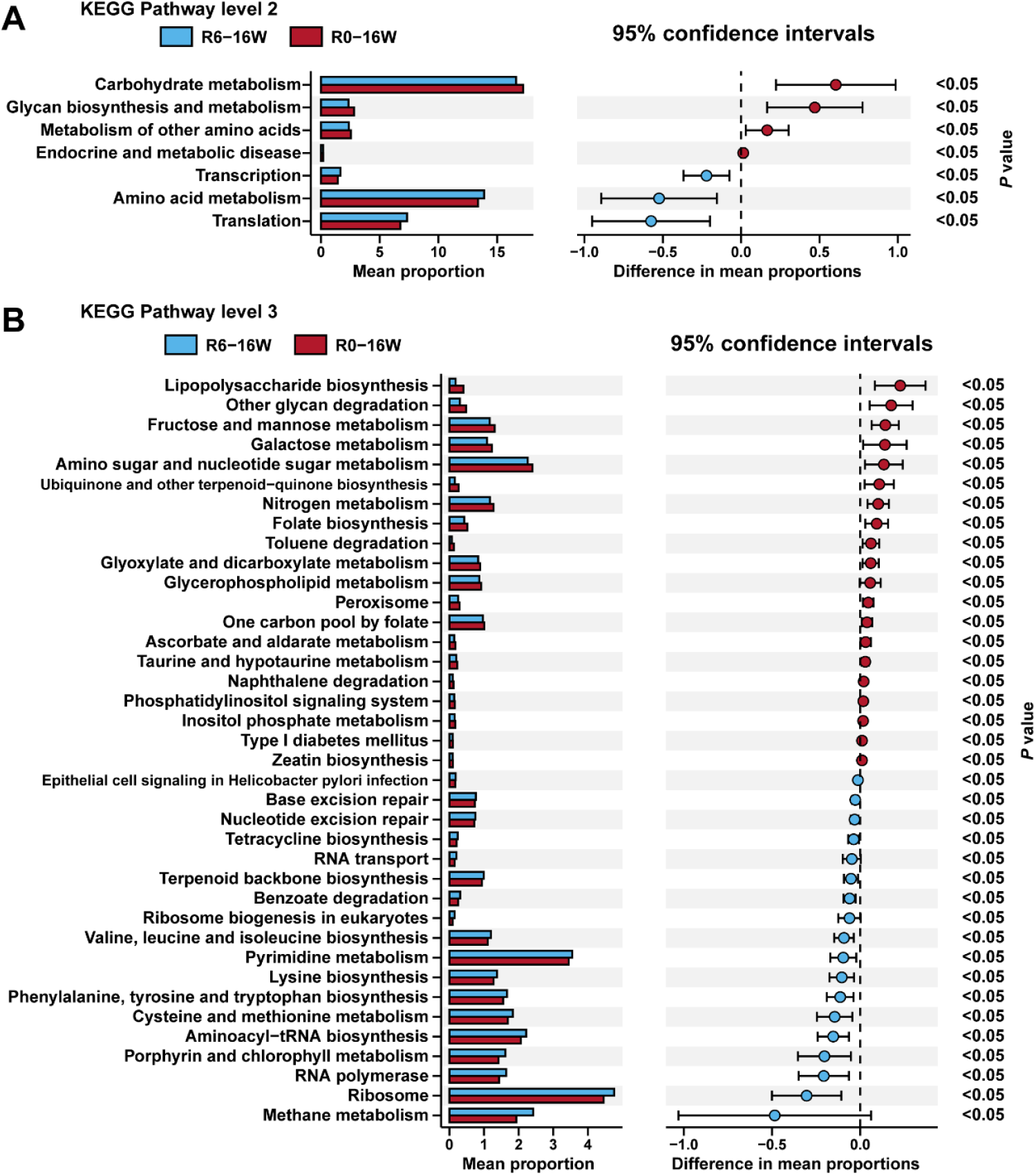
Effects of riboflavin deficiency on gut microbiota functions predicted by PICRUSt2. The differences in gut microbiota functional profiles between R0-16W rats and R6-16W rats at KEGG pathway level 2 (**A**) and KEGG pathway level 3 (**B**).

**Figure 7.**
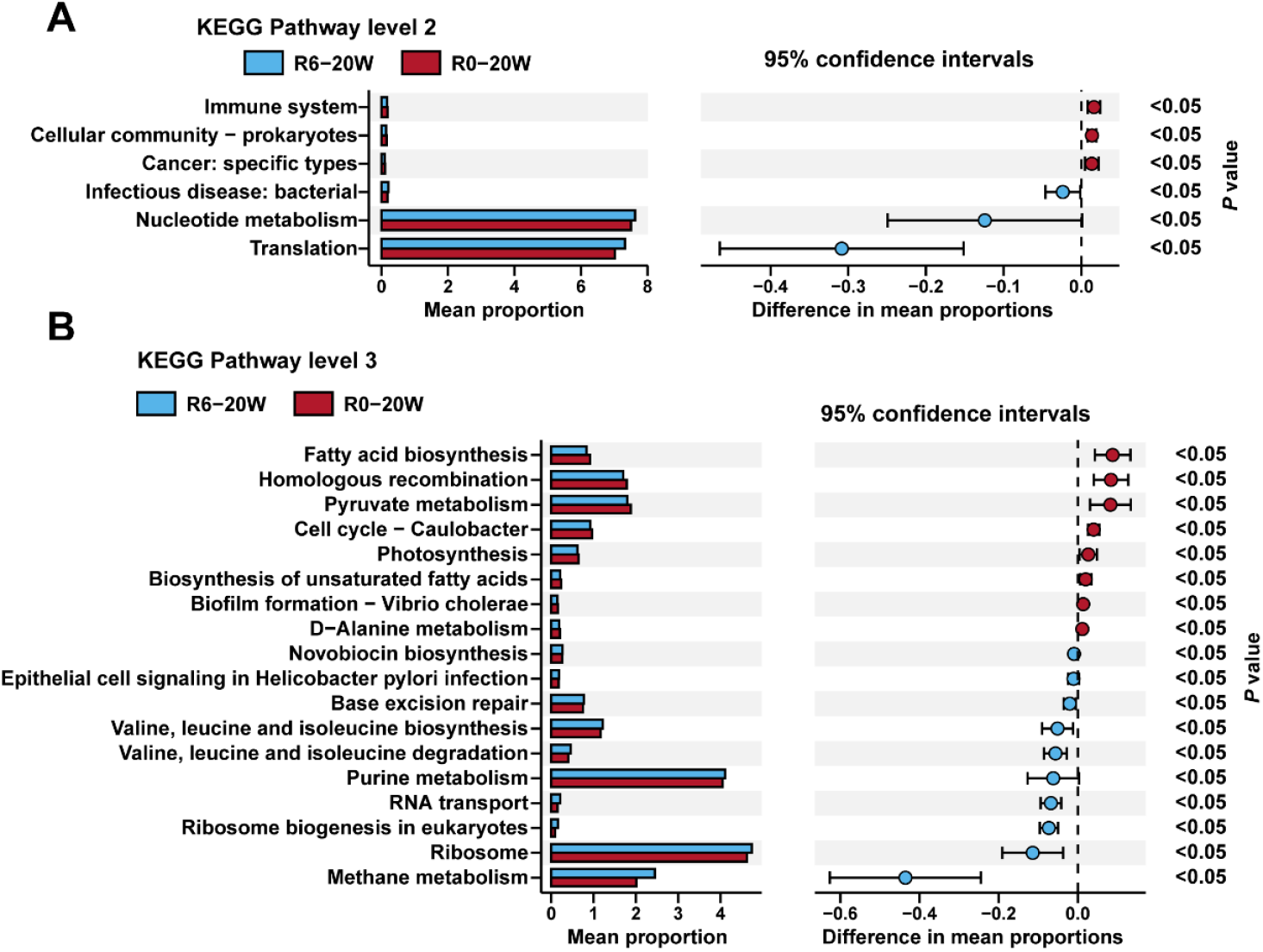
Effects of riboflavin replenishment on gut microbiota functions predicted by PICRUSt2. The differences in gut microbiota functional profiles between R0-20W rats and R6-20W rats at KEGG pathway level 2 (**A**) and KEGG pathway level 3 (**B**).

## Discussion

In humans, riboflavin deficiency leads to ariboflavinosis, characterized by symptoms such as sore throat, glossitis, angular stomatitis, conjunctivitis, and seborrheic dermatitis, among others^[8, 9]^. However, the precise mechanisms underlying riboflavin deficiency and the development of ariboflavinosis have not been fully elucidated. In this study, we confirmed that riboflavin deficiency causes ariboflavinosis (manifesting as angular stomatitis and conjunctivitis) in rats and revealed that it is closely associated with alterations in gut microbiota composition.

The taxonomic chain Proteobacteria-Gammaproteobacteria-Enterobacteriales-Enterobacteriaceae-*Escherichia-Shigella* is associated with ariboflavinosis. Gut dysbiosis, characterized by a marked expansion of genus *Escherichia-Shigella*, is a hallmark of rats with ariboflavinosis and may serve as a promising diagnostic biomarker and therapeutic target for ariboflavinosis. Studies have shown that in ulcerative colitis, compared with non-inflamed mucosae, Proteobacteria ((notably *Escherichia-Shigella*) were increased in inflamed mucosae. The microbial dysbiosis, primarily characterized by an increased abundance of the *Escherichia-Shigella* genus, is a feature of the inflamed mucosae in ulcerative colitis patients^[20]^. The significant gut microbiota dysbiosis observed in tuberculous meningitis patients was associated with markedly high proportions of *Escherichia-Shigella* species as well as increased blood levels of tumor necrosis factor alpha (TNF-α) and interleukin 6 (IL-6) ^[21]^. Amyloidosis patients showed higher abundance of *Escherichia-Shigella*. A positive correlation was observed between pro-inflammatory cytokines IL-1β, NLRP3, and CXCL2 with abundance of the inflammatory bacteria taxon *Escherichia-Shigella*^[22]^. High ammonia induced lung tissue inflammation through increasing the proportion of *Escherichia-Shigella*, activating the NLRP3 inflammasome, and promoting IL-1β release^[23]^. These findings suggest that further studies are warranted to investigate the potential contribution of *Escherichia-Shigella* in the pathogenesis of ariboflavinosis.

The gut microbiota inhabits the gastrointestinal tract and exerts both local and systemic effects. The interaction between the gut microbiota and the host occurs through microbial metabolites, hormonal intermediates, and immunologic messengers ^[24]^. Riboflavin deficiency leads to increased lipopolysaccharide (LPS) biosynthesis by bacteria. Microbes residing in the host’s gastrointestinal tract release signaling byproducts from their cell wall, such as LPS, which can act both locally and, after crossing the gut barrier and entering the circulation, systemically^[25]^. The gut microbiota controls the development of the immune system by setting systemic threshold for immune activation. LPS from gut bacteria have been shown to elicit both systemic pro-inflammatory and immunomodulatory responses. It has been established that ingestion of a high-fat diet increases the blood levels of LPS from Gram-negative bacteria in the gut. Experimental evidence shows that LPS is involved in the transition of macrophages from the M2 to the M1 phenotype^[26]^. We speculate that the bacteria implicated in ariboflavinosis may act through LPS, a bacterial metabolite, which requires further investigation.

## Conclusions

The present study demonstrates that riboflavin deficiency induces ariboflavinosis in rats. Riboflavin deficiency modulates the composition of the gut microbiota. Functional prediction suggests that lipopolysaccharide biosynthesis may be a potential mechanism. Our findings provide new insight into riboflavin deficiency and ariboflavinosis. However, further investigations will be necessary to establish the precise causal relationship.

## Abbreviations

FLASH: Fast Length Adjustment of SHort Reads
g: Gram
nM: Nanomoles per liter (nmol/L)
OTUs: Operational taxonomic units
PCoA: Principal coordinates analysis
PICRUSt2: Phylogenetic Investigation of Communities by Reconstruction of Unobserved States 2
QIIME2: Quantitative Insights Ioto Microbial Ecology 2
R6: Riboflavin 6 mg/kg
R0: Riboflavin 0 mg/kg
W: Week.

## Declarations

### Ethics approval and consent to participate

The experimental procedures administered on animals were approved by the Institutional Animal Care and Use Committee of Shantou University.

### Consent for publication

Not applicable.

### Availability of data and materials

All data are available in the manuscript or upon request to the authors

### Competing interests

The authors declare that they have no competing interests.

### Funding

This work was supported by grants from the Science and Technology Planning Project of Guangdong Province of China (210713176903477 and 210715116901229);the Project of Administration of Traditional Chinese Medicine of Guangdong Province of China (20241295 and 20251357); Guangdong Basic and Applied Basic Research Foundation Enterprise Joint Fund (2023A1515220058); the Medical Scientific Research Foundation of Guangdong Province (A2024738 and B2025516).

### Author contributions

FP planned the studies. WZ, PPW, WQS and HZ performed the experiments. WZ held the responsibility for all data integrity and data analysis. FP conducted the entire research project. All authors have read and approved the final manuscript.

## Acknowledgments

We are grateful for the assistance provided by the Central Laboratory at Shantou Central Hospital, particularly to Prof. Ze-peng Du, who helped us gain access to an excellent instrumental platform.

## Conflict of interest

The authors declare no conflict of interest.

## Code availability

Not applicable.

## Figure legends

**Supplemental Figure 1.**
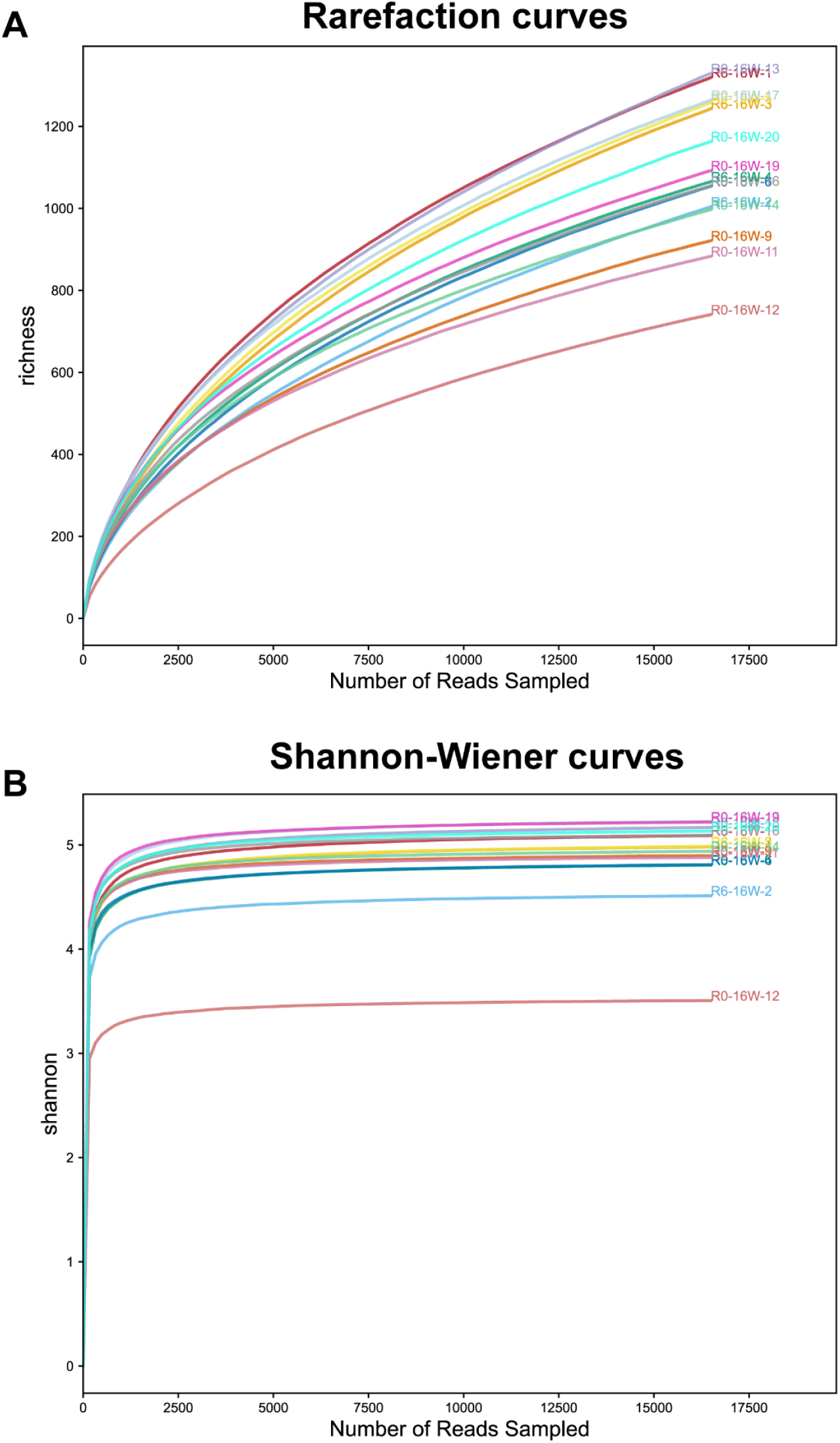
**Rarefaction curves (A) and Shannon-Wiener curves (B).**

**Supplemental Figure 2.**
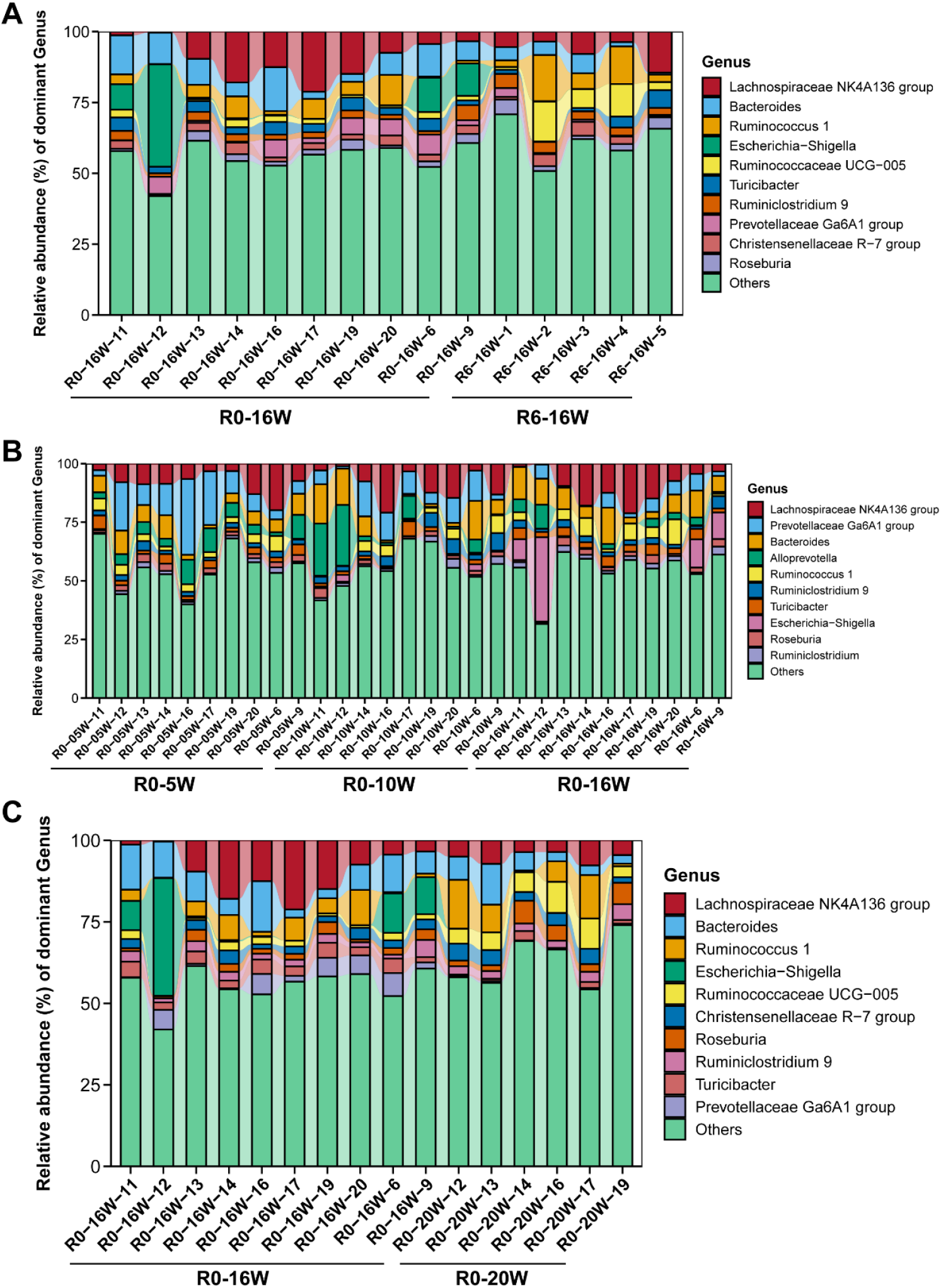
Riboflavin deficiency modulates gut microbiota composition in rats. (**A-C**) Gut microbiota composition at the genus level. Each column represents the relative abundance of bacteria for one rat. *n* = 5-10/group (angular stomatitis and conjunctivitis in R0-16W group).

## Notes

### Competing Interest Statement

The authors have declared no competing interest.

